# A composite method to infer drug resistance with mixed genomic data

**DOI:** 10.1101/2020.07.30.194266

**Authors:** Gargi Datta, Nabeeh A Hasan, Michael Strong, Sonia M Leach

**Author notes:** Authors contributed equally in a senior role (Michael Strong, Sonia M Leach). Email addresses (Gargi Datta), (Nabeeh Hasan), (Michael Strong), (Sonia Leach).

## Abstract

**Background:** The increasing incidence of drug resistance in tuberculosis and other infectious diseases poses an escalating cause for concern, emphasizing the urgent need to devise robust computational and molecular methods identify drug resistant strains. Although machine learning-based approaches using whole-genome sequence data can facilitate the inference of drug resistance, current implementations do not optimally take advantage of information in public databases and are not robust for small sample sizes and mixed attribute types.

**Results:** In this paper we introduce the Composite MetaDistance method, an approach for feature selection and classification of high-dimensional, unbalanced datasets with mixed attribute features from various data sources. We introduce a mixed-attribute, multi-view distance function to calculate distances between samples, with optimal handling of nominal features and different feature views. We also introduce a novel feature set for drug resistance prediction in *Mycobacterium tuberculosis*, using data from diverse sources. We compare the performance of Composite MetaDistance to multiple machine learning algorithms for *Mycobacterium tuberculosis* drug resistance prediction for three drugs. Composite MetaDistance consistently outperforms existing algorithms for small sample training sets, and performs as well as other algorithms for training sets with larger sample sizes.

**Conclusion:** The feature set formulation introduced in this paper is utilizes mutational and publicly available information for each gene, and is much richer than ever devised previously. The prediction algorithm, Composite MetaDistance, is sample size agnostic and robust especially given small sample sizes. Proper handling of nominal features improves performance even with a very small number of nominal features. We expect Composite MetaDistance to be even more robust for datasets with a higher percentage of nominal features. The algorithm is application independent and can be used for any mixed attribute dataset.

## Background

In recent years, the emergence of antibiotic drug resistance has been a growing concern to public health, with an unprecedented number of resistant microorganisms and an ever-increasing cache of drugs to which microorganisms are resistant [1–3]. Diseases like tuberculosis (TB) are adversely affected by the emergence of multiple drug-resistant (MDR) and extensively drug resistant (XDR) strains [4]. The World Health Organization (WHO) prescribed treatment for drug-susceptible TB is a six-month regimen of the first line drugs: Rifampin, Isoniazid, Ethambutol and Pyrazinamide [5]. Drug-resistant cases are prescribed additional drugs with increased toxicity and extensive side effects, like the second-line drug classes of Fluorquinolones and Aminogylcosides [5]. Current WHO standards of drug susceptibility testing include phenotypic susceptibility testing, which involve growing cell-cultures of the bacteria in the presence of a particular drug, in order to determine if the strain is resistant or susceptible to that drug [4, 6, 7]. This can take several weeks to months in some cases, resulting in a lag in optimal drug selection and contributes to the potential transmission of drug-resistant strains [4]. Phenotypic susceptibility tests are reliable for some drugs, like Isoniazid and Rifampin, but they tend to be difficult to conduct and have low sensitivity for others drugs including Ethambutol and Pyrazinamide [7]. Recently, molecular drug-susceptibility testing methods have gained momentum, but many of these assays only test for a few well-known mutations, and in some cases can have low-sensitivity for certain compounds [4, 7, 8]. Molecular methods are especially unreliable for smear-negative, culture positive sputum [7]. Understanding the mechanisms for drug resistance and developing methods to accurately predict resistance of new disease strains of global importance is vital to optimal treatment [1, 4, 8]. Since the majority of cases of drug resistance in *M. tb* occur due to mutations in genes that encode target proteins or drug activating enzymes and their promoter regions, the problem of drug resistance prediction can be broken down into a per-gene per-drug prediction problem [8, 9]. This can be defined as whether a mutation in a gene that a drug binds to confers resistance to that drug. For instance, the majority of cases of resistance to Rifampin, Isoniazid and Pyrazinamide can be attributed to mutations in their respective primary gene targets and activating enzymes, *rpoB*, *katG*, and *pncA* respectively (Table 2) [5, 9]. Thus, predicting drug-resistance in *M. tb* reduces to a per-gene per-drug resistance prediction task [6, 8, 10, 12].

The introduction of whole genome sequencing (WGS) using next generation sequencing methods has paved the path for quick identification of resistance-causing mutations in target genes, and has led to the application of machine learning (ML) approaches based on sequence information to rapidly predict drug resistance of novel strains [6]. Current WGS-based ML approaches use a single type of feature per gene to predict resistance to a particular drug [8, 10, 12]. Niehaus *et al.* used the presence or absence of a single nucleotide variation (SNV) in resistance-conferring genes in *M. tb* to classify strains as resistant or susceptible, whereas Walker *et al.* examined specific mutations that are known to impart resistance to first-line and second-line TB drugs, and predicted resistance to drugs by evaluating the presence and absence of those specific mutations [8, 10]. Yu *et al.* applied Delaunay triangulation to encode genotypic information on protein structures to predict resistance to HIV drugs [11]. While focusing on a single feature type such as sequence features or structure features simplifies the prediction process, these features fail to incorporate the wealth of publicly available information that can be derived from sequence data. This research tackles the problem by defining a robust and complete feature set with different types of features derived from genomic information and publicly available databases, including whether one amino acid substitution has a larger effect on the protein than another, whether mutations are located in a region of the protein known to have high resistance to the particular antibiotic, or whether the strain is similar to a highly virulent or know resistant strain [13]. Such a diverse and robust feature set is the first such feature set in drug resistance prediction, especially in *M. tb*. In addition, the feature set can be extended further to include additional information, such as type of amino acid change or binding affinity.

The introduction and application of complex feature sets for the inference of drug resistance presents additional challenges to machine learning algorithms, including increased dimensionality, redundancy among features and the presence of a mix of continuous, ordinal and nominal attribute types. In addition, most real-world datasets are extremely unbalanced. Finally, while a few of the first-line drugs have publicly available susceptibility information for thousands of strains, the second- and third-line drugs for tuberculosis lack such wealth of information. Furthermore, to model an isolated outbreak of *M. tb*, or another organism without a wide range of sequences of strains available, robust methods are needed to prevent overfitting for small sample sizes. We present a method that addresses each of these issues, by combining an approach that simultaneously performs classification and feature selection with a distance metric sensitive to mixed attribute types, using a robust heterogeneous feature set based on sequence, structure, amino acid substitutions, functional domains, location and phylogeny information. Our method outperforms existing algorithms for small sample sizes. We first motivate the approach and provide an overview (Fig 1, 2, 3; More information in methods).

**Figure 1.**
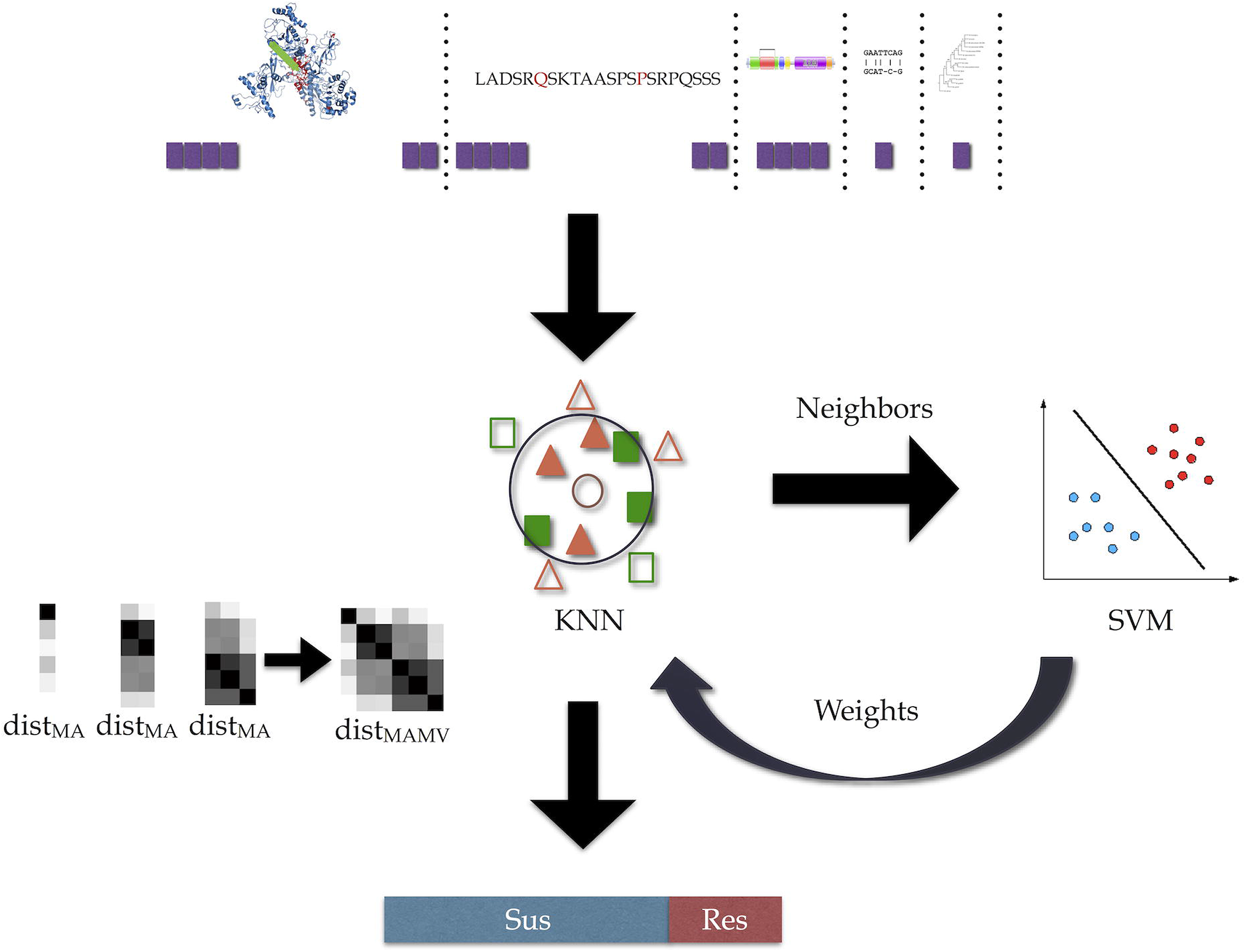
Overview of Composite MetaDistance. Figure 1 shows the schematic of Composite MetaDistance. The base algorithm, MetaDistance uses a k-nearest neighbor algorithm for classification and an SVM formulation for feature importance calculations. Composite MetaDistance implements a mixed-attribute multi-view distance function in conjunction with MetaDistance to classify mixed attribute data from multiple data sources.

**Figure 2.**
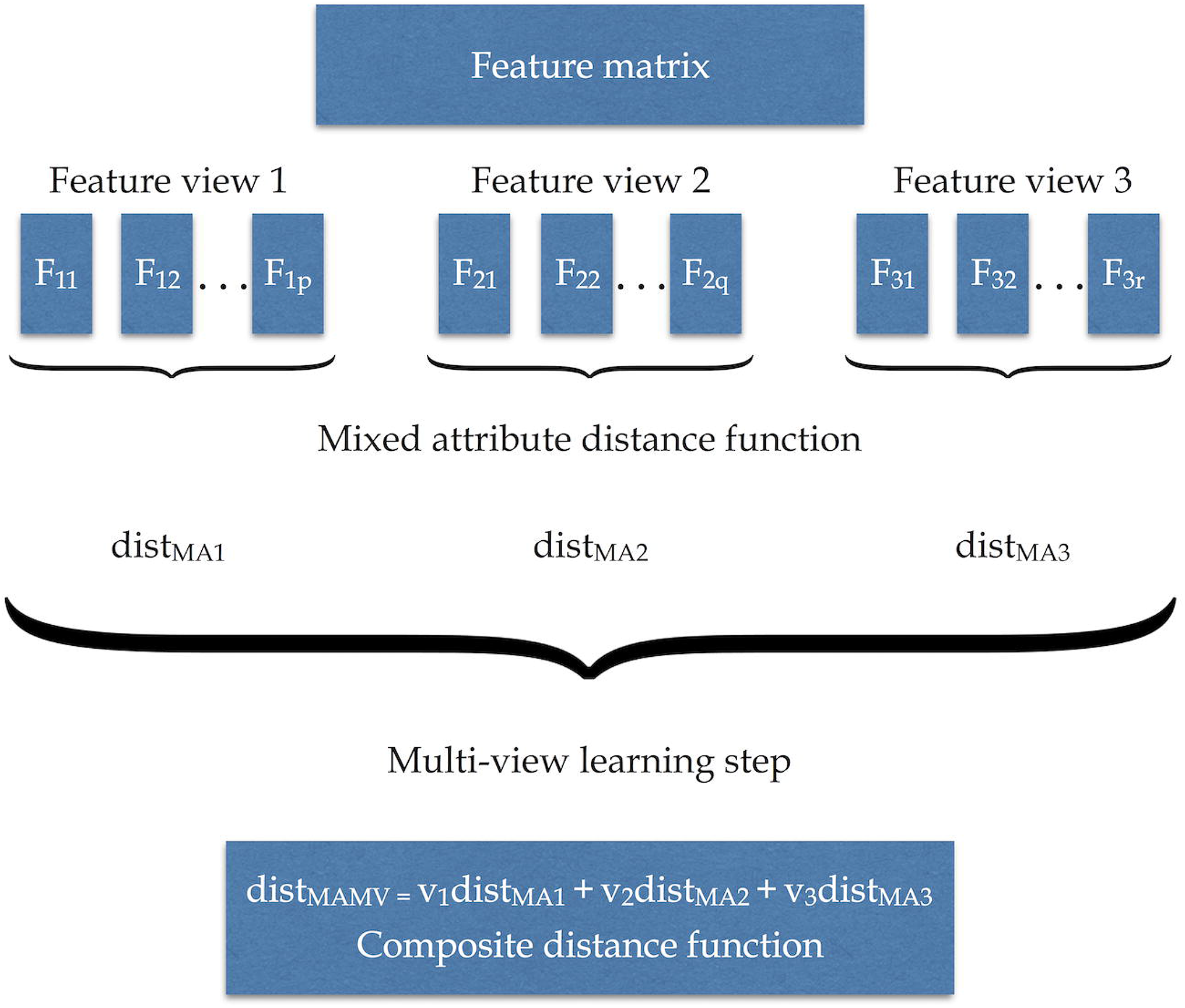
Overview of Composite Distance Function. A mixed attribute distance function (dist_MA_) is used to calculate distances between samples for each feature in a view and combine the distances per view. The final distance between two samples is then calculated by a linear combination of the dist_MA_ for each view (multi-view learning step). The final distance function is called the mixed-attribute multi-view distance function (dist_MAMV_), or the composite distance function.

**Figure 3.**
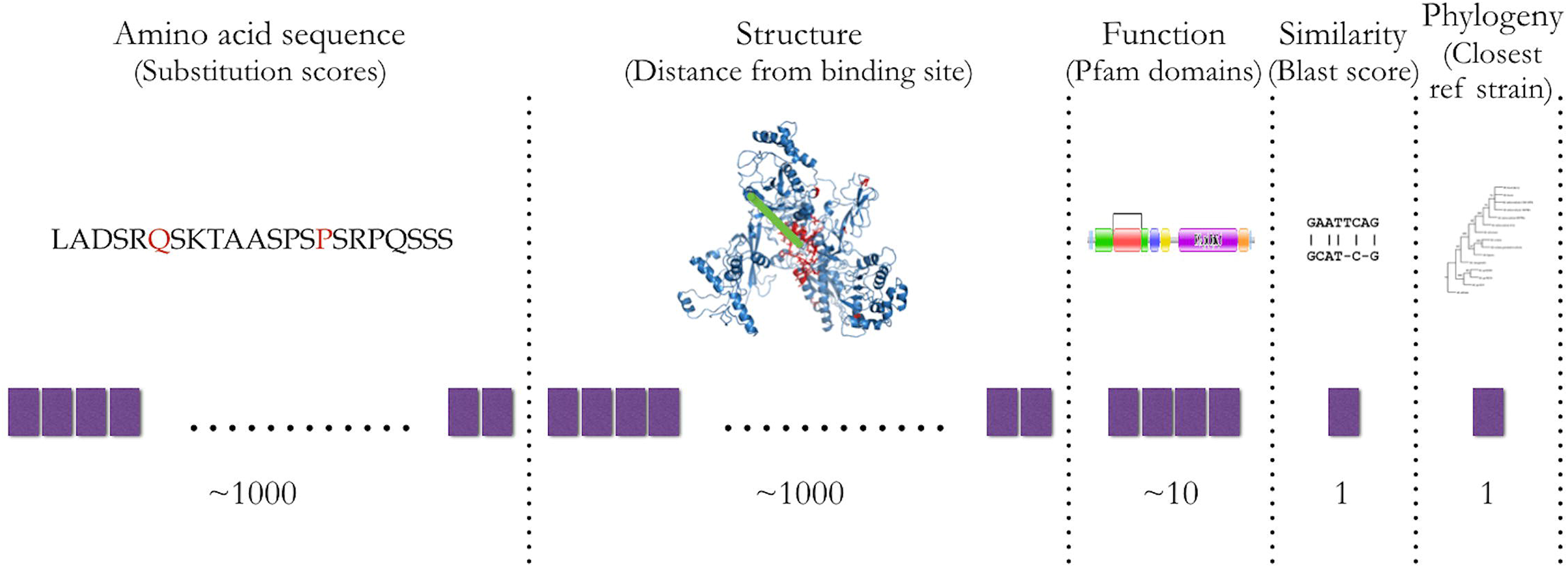
Feature set schematic. The feature set contains features from different data sources. There are ~1000 structural and amino acid-based features, 1-10 function based features, 1 similarity based feature and 1 phylogeny based feature. These are all derived from the sequence information and publicly available data sources.

### Mixed Attribute Types

Our feature set includes variables with mixed attribute types, including a continuous variable representing the distance of an amino acid to the known drug binding site, an ordinal variable representing the level of tolerance for a particular amino acid substitution, and a nominal variable representing the closest reference strain. Most modern classification and clustering algorithms are designed to work well with continuous data, and relatively few are equipped to handle mixed data in a proper manner [14–17]. Tree ensemble methods, such as random forests, can grow trees for different variable types separately before performing an ensemble variable selection [15, 18]. Although useful, random forests show selection bias favoring variables with more values, and variable selection is affected by the scale of measurement of predictor variables, thus making them unreliable for mixed data [15, 18]. Nominal features are especially tricky to handle for most traditional algorithms. Distance metrics are usually designed for continuous (and ordinal) data and do not work with nominal variables. While continuous and ordinal variables have an intrinsic order, there is a lack of order among the individual attribute values for categorical or nominal variables, thus making it difficult to define a distance metric [14, 17]. Most uses of nominal variables assume an initial enumeration of the distinct categories and are treated by the learner as a continuous or ordinal variable. For example, any algorithm with a Manhattan or Euclidean distance metric (KNN or SVM for example) treats nominal variables as continuous. The magnitude of the distance between two items is thus dependent on their proximity in the ordering imposed by the initial enumeration, rather than capturing the fact that they are dissimilar or not. Thus, choosing an appropriate distance metric for high-dimensional mixed data is an important factor in correctly classifying and clustering datasets with mixed attribute types [14, 19]. To address this problem, we use an extension of Gower’s General Similarity Coefficient as a distance metric for mixed data [16, 19]. Here, the distance between two samples for a feature is a range-normalized difference between the values if the feature is continuous or ordinal. If the feature is nominal, the distance is a binary representation indicating if the categories are the same or not [16, 17, 19]. The final distance between two vectors of mixed attributes is then the sum of the per-attribute distances. More details about the use of a mixed attribute distance are given in methods.

### Multi-view learning

Our feature sets for drug resistance prediction originate from different feature sources (or views, as in database theory). The view provided from structural information captures the 3-D distance of a mutation to the known drug-binding site, while the view provided from sequence in-formation captures the impact of an amino acid mutation on the sequence. Functional information captures the presence of mutations in specific functional domains, phylogenetic information shows whether a strain is closely related to a highly virulent or resistant reference strain and BLAST score captures the similarity of the strain to the reference strain H37Rv. Classical machine learning algorithms typically concatenate features from all views into a single large feature vector used for learning and thus may be subject to overfitting [20]. Since some views have thousands of features, and some have one feature, this causes imbalance in how a view is weighted and considered by the algorithm. In addition, different views may benefit from using different similarity measures, which is not accounted for by classical machine learning algorithms [21]. Multi-view learning focuses on each view individually before jointly optimizing them for the final classification [20, 22]. Multiple kernel learning is a class of algorithms for multi-view learning, where each kernel captures the notion of similarity most appropriate, and kernels are combined either linearly or non-linearly [20–22]. The weights used for the kernel combination signify the relevance of each view for predicting drug resistance. We use the principles of multiple kernel learning for drug-resistance prediction by first computing the distance between two strains per feature view, using the distance measure for mixed attribute types described above. The composite distance between two strains is then calculated as a weighted sum of the view-specific distance matrices. This composite distance is used by a learner for training and testing.

### Uninformative features and unbalanced data with small sample sizes

Feature sets proposed for drug resistance tend to have highly redundant and uninformative features [1, 8, 10]. For example, the feature set may include all amino acid positions yet there may be no mutations recorded in a particular amino acid for a given dataset, thus making the structure-based and sequence-based features for that amino acid uninformative. Both mentioned properties are reflected in our data, where the same uninformative feature may be repeated in the structure and amino acid-based feature views, and our datasets are extremely imbalanced. Thus, it is critical to have a classification algorithm that can handle high-dimensional, unbalanced datasets with redundant, mixed attribute types from multiple feature views. In addition, a large sample size is not always available. Until recently, *M. tb* studies had only a few hundred samples. Publicly available data for some second-line drugs for tuberculosis still have a very small number of strains (~100s) with phenotypic susceptibility data available, thus making this a small-sample classification problem. Most model-based machine-learning techniques, such as support vector machines (SVM) and logistic regression, work well with high-dimensional data, but their performance decreases with small sample size [24]. Instance-based methods like k-nearest neighbor (KNN) on the other hand work well with small samples, but their accuracy decreases with an increase in uninformative and redundant features [24]. To address these difficulties, we leverage an approach originally designed for metagenomics projects, called MetaDistance [24]. MetaDistance combines model- and instance-based methods to simultaneously perform classification and feature selection. Our method uses MetaDistance in conjunction with the modified Gower’s similarity coefficient and multiple kernel learning principles for robust prediction of drug resistance in TB and other bacteria [21, 24, 25]. Fig. 1 shows a schematic of the overall method, which we named *Composite MetaDistance*.

## Methods

For the drug-resistance prediction task, we assume that mutations in the drug target gene affect the susceptibility or resistance to the drug of a given *M. tb* strain. For each strain, a set of features is created based on sequence, structure, amino acid substitutions, functional domains, location and phylogeny information, where each information source contributes a view of the data comprised of a mix of continuous, ordinal and nominal attributes. Thus, the input data for a given sample (strain) *i* consists of a vector ***x***_*i*_ of features and the output data *y* consists of a categorical variable indicating susceptibility or resistance (bold indicates vector). Our method extends an existing classification algorithm by offering an alternative distance measure to correctly handle mixed attribute types and to appropriately integrate multiple heterogeneous input data (views) into a single distance function. Here we describe the base algorithm, outline our extensions, and describe the input data and feature sets.

### Feature set per gene

The per-gene feature set includes features from multiple views. These views include features based on structure, amino acid substitutions, functional domains, similarity to the reference strain, and phylogeny (Fig. 3, Table 1).

**Table 1.** Number of features per datatype for each gene

| Gene | # Continuous | #Ordinal | #Nominal |
| --- | --- | --- | --- |
| *katG* | 741 | 740 | 4 |
| *pncA* | 187 | 186 | 2 |
| *rpoB* | 1173 | 1172 | 7 |

#### Amino acid-based features

Amino acid-based features capture the observation that some amino acid substitutions are more tolerated by the protein structure than others. If a gene has n amino acids, there are n amino acid-based features for each strain for that gene. For each *M. tb* strain, the value for each feature in the amino acid-based features is:

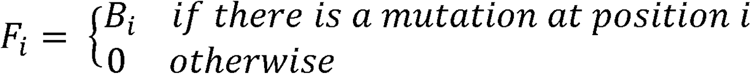

where B_i_ denotes the BLOSUM62 score of the amino acid substitution position i [43]. Since the BLOSUM62 values are integers between −4 to 11, the amino acid-based features are all ordinal features.

#### Structure-based features

Structure-based features are motivated by the observation that mutations in the binding site for a drug are more likely to affect susceptibility and use the 3-D protein-drug structure to measure amino acid proximity to the binding site. If a gene has n amino acids, there are n structure-based features for each strain for that gene. For each *M. tb* strain, the value for each feature in the structure-based features is:

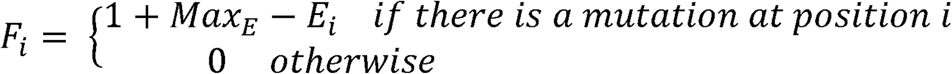

where E_i_ denotes the Euclidean distance of amino acid position i from the center of the binding site for the particular drug to the gene. The center of the binding site for a drug for the gene is the average of the x, y and z positions of the amino acids in the known binding site for that drug to the protein. Max_E_ is the Euclidean distance of the amino acid position farthest away from the center of the binding site. The structure-based features are all continuous features.

#### Function-based features

The function-based features detect whether a mutation exists in a particular functional domain, since mutations may inhibit that domain and affect the protein’s function. The number of function-based features are determined by the number of functional domains for the gene reported in PFAM version 29.0 [44]. For a gene with p PFAM domains, there are p functional features. These are binary features:

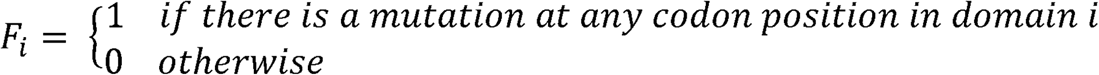

for each domain i in PFAM for the protein. The function-based features are all nominal features.

#### Similarity-based feature

The similarity-based feature is a single feature that captures how divergent is the strain from the reference. This single continuous feature is defined as

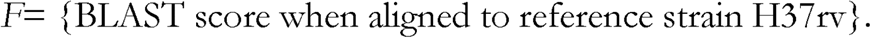

The BLAST score used here is the bit score calculated by BLAST. H37Rv is the standard reference strain used for *M. tb* sequence analysis and the datasets were aligned to this strain in the mutation identification and annotation pipeline. The BLAST scores were calculated using BLAST 2.2.28+ [45].

#### Phylogeny-based feature

The phylogeny-based feature builds upon the inference that since drug resistance and susceptibility are known for reference strains, knowing the closest reference strain could prove informative [13]. The set of reference strains include 10 strains representing the major groups in the *M. tb* complex. All sequences for each gene were collapsed into unique haplotypes using FaBox DNACollapser to reduce redundancy [46]. We constructed rooted phylogenetic trees based on the observed distances, using the neighbor-joining method as implemented in SeaView 4.4.0 [47]. Bootstrap support was calculated using 100 replicates. The closest reference strains were inferred from these phylogenetic trees. The phylogeny-based feature is a nominal feature with 10 possible values representing the closest reference strain.

#### Creating the per-gene feature set from individual feature views

The feature set per-gene is a concatenation of all the individual features for that gene (Fig. 1). The amino acid substitution-based features are created as the first feature view for genes that transcribe to proteins. If the protein structure and binding site for a drug are known, the second set of features (second view) is the structure-based features. If PFAM domain information is available for a gene, the function-based features are appended as the third feature view. Then, the phylogeny-based feature is appended to the feature set, and finally the similarity-based feature is added to the feature set. It is important to note that the sequence of views is not important in the concatenation, as long as it is consistent.

### Composite MetaDistance

For the per-gene per-drug resistance prediction task, it is assumed that mutations in the drug target gene or its promoter region affect the susceptibility or resistance to the drug of a given *M. tb* strain. For each strain, a set of features is created based on sequence, structure, amino acid substitutions, functional domains, location and phylogeny information, where each information source contributes a view of the data comprised of a mix of continuous, ordinal and nominal attributes. Thus, the input data for a given sample (strain) i consists of a vector ***x***_***i***_ of features and the output data y consists of a categorical variable indicating susceptibility or resistance (bold indicates vector). Prediction errors, learned weights for each feature and neighborhood scores for each strain in the input data for the two target classes are also provided. The Composite MetaDistance method extends an existing classification algorithm, MetaDistance, by offering an alternative distance measure to correctly handle mixed attribute types and to appropriately integrate multiple heterogeneous input data (views) into a single distance function [24].

#### Base Algorithm - MetaDistance

MetaDistance, the base classification algorithm, combines a k-nearest neighbor (KNN) with a support vector machine formulation (SVM) to simultaneously select features and classify samples, by jointly maximizing the interclass distance and minimizing the intraclass distance between samples [24]. The KNN formulation in MetaDistance is a modification of the basic KNN (mod-KNN), and deals with unbalanced data. To predict the class of a sample, MetaDistance calculates the k closest neighbors to a sample in each class separately, and assigns to the sample the class with the closest mean-neighborhood score. The authors of MetaDistance determined k by cross validation with the smallest prediction error [24]. This modified KNN algorithm is also used in Composite MetaDistance.

The (simple attribute) distance between two samples *i* and *j* is denoted by:

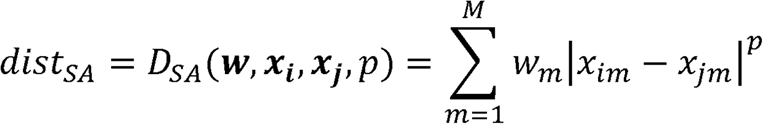

where ***x***_***i***_ and ***x***_***j***_ are feature vectors for samples *i* and *j* respectively, *w*_*m*_ ≥ 0 are non-negative weights for *m* ∈ [1, *M*] features, and *p* is a positive free parameter that can be initialized to different values for different distances, e.g. *p*=1 for city block, *p*=2 for Euclidean and so on [24]. *w*_*m*_ and *p* are initialized to 1.0 by default in the algorithms. To test a new sample, we first compute the average distance per class among the k-nearest neighbors assigned that class, and then we return the class with the minimal average distance vote.

During training of MetaDistance, the KNN uses *dist*_*SA*_ to define the neighborhood for each sample, for each class, and then passes that neighborhood to a quadratic SVM formulation to optimize the feature weights **w** [24]. In addition to the standard SVM constraints used to optimize the margin, and therefore maximize interclass distance, the MetaDistance SVM formulation also includes constraints to minimize intraclass distances among the neighborhood defined by the KNN. The SVM uses a standard conjugate gradient method to estimate the weights (**w**) [24, 26]. Feature importance can be determined from the weights. The estimated weights are then passed back to the KNN, which uses the weights with *dist*_*SA*_ to classify test samples [24].

#### Composite Distance Function

For drug resistance prediction with mixed attribute types from different feature views, we created a composite distance function for mixed attributes based on Gower’s similarity, in addition to a multi-view kernel step, to calculate the distance between two samples [19–21, 25]. Fig. 2 shows a schematic of the overall method.

Our approach first builds on the basic MetaDistance by substituting a mixed attribute distance function (*dist*_*MA*_) i in the KNN:

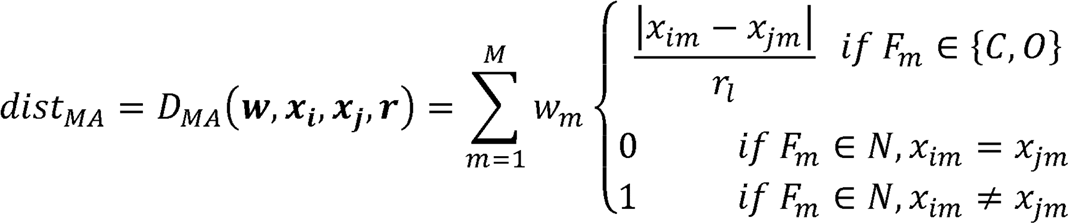

where ***x***_***i***_ and ***x***_***j***_are feature vectors for samples *i* and *j* respectively, and *w*_*m*_ ≥ 0 are non-negative weights for *m* ∈ [1, *M*] features. *r*_*m*_ is the range of values for feature *F*_*m*_ for continuous and ordinal features, and is calculated as:

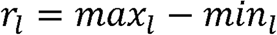

*C*, *O* and *N* denote the continuous, ordinal, and nominal feature spaces respectively. Note that regardless of attribute type, the per-attribute distance term is a value between 0.0 and 1.0.

The distance function *dist*_*MA*_ accounts for mixed attribute types, but does not address the problem of multiple feature views. To properly combine distances between features of different feature sets, we introduce a multi-view learning step to our method, thus creating a new mixed-attribute multi-view distance function (*dist*_*M AMV*_) as a linear combination of all the feature views:

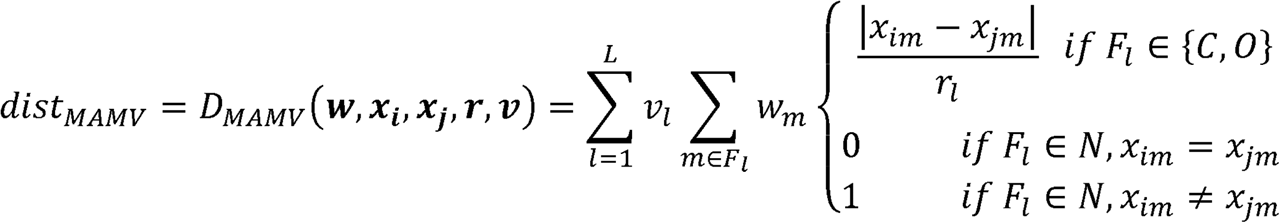

where *l* ∈ [1, *L*] denotes each feature view and *m* ∈ *F*_*l*_ denotes all the features in view *l*. The multi-view coefficient *v*_*l*_ is the weight for view *l*, which is used along with the feature weights *w*_*m*_ to determine the closest neighbors for a sample.

Our method, Composite MetaDistance, thus extends MetaDistance by substituting the distance function used by the KNN to the mixed-attribute multi-view distance function (*dist*_*MAMV*_) to find the *k* nearest neighbors for each sample. We determine *k* as the value with the lowest training error, considering *k* between 1 and 10. Learning proceeds as in MetaDistance to optimize the feature weights **w**, which are then used by the KNN to classify test samples.

### Feature and view coefficients

#### Multi-view coefficients

Many strategies could be chosen to normalize the distances for each feature view. A simple alternative is to set the multi-view coefficients v_l_ to the inverse of the number of features in each view l. Thus, the final distance represents the average distance per view. This removes the bias towards views with more features and each view is considered equally in the final sum.

#### Feature weights

For the learning step, the feature weights (*w*) are initialized to 1.0, and then the quadratic SVM is applied to learn the weights. As an alternative, we performed an individual feature analysis with a random forest classifier (RandomForestClassifier in scikit-learn 0.16.1, Python 2.7.10) on our training data to understand how well each feature performs as an independent predictor. The feature weights (*w*) for each feature *m* were then initialized to the area under the receiver-operator curve (AUC) per feature from the individual feature analysis:

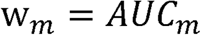

#### *M. tuberculosis* data

The first step in any machine learning problem is to find adequate data to train on. Unfortunately, for most diseases, there is no centralized data bank to store and download thousands of sequences easily. For organisms like Nontuberculous Mycobacteria (NTM), no more than a few hundred genomes have been sequenced and are publicly available. Recently, with the rapid development and cost-effectiveness of whole genome sequencing, initiatives have been made to sequence thousands of strains of major infectious agents, such as *M. tb* and HIV [10, 51]. Walker *et al.*, in late 2015, published a study with sequence data from 3,651 *M. tb* strains, from UK, Sierra Leone, South Africa, Germany and Uzbekistan, representing all seven clades of TB [10]. These strains were tested by Walker *et al.* for phenotypic susceptibility to the first-line drugs [10]. This dataset is unique in the observation that less than 20% of *M. tb* strains globally are antibiotic resistance is reflected in the imbalance of the dataset [5]. The whole genome sequences for these strains were deposited in NCBI and EBI. This was the first such large-scale study for tuberculosis, and enabled us to download the sequences for 3,650 of those strains for the machine learning analysis [10]. The raw sequence files downloaded for the 3,650 strains were run through an automated mutation identification and annotation pipeline outlined to get mutations in resistance-conferring genes (Table 2) [33]. Resistance to drugs in *M. tb* is usually due to mutations in the genes that bind or activate drugs [9]. Features for drug resistance prediction for each gene were then derived using those mutations along with publicly available data. Drug resistance to Rifampin, Isoniazid, and Pyrazinamide was predicted using feature sets for the appropriate drug target genes (Table 2). We predicted resistance of strains to one or more of three first line drugs: Isoniazid, Pyrazinamide and Rifampin by creating feature sets for mutations in their primary resistance associated genes, *katG*, *pncA* and *rpoB* respectively [5]. *rpoB* is known to account for 95% of resistance to the drug Rifampin [27]. Similarly, mutations in *katG* and *pncA* account for the majority of resistance to Isoniazid and Pyrazinamide respectively [28, 29]. These per-gene feature sets were then used to predict resistance to the respective drug. This dataset of 3,650 strains will henceforth be referred to as the Global Dataset.

**Table 2.** Number of *M. tb* strains per gene in the Global dataset

| Drug | Gene | Training set | Susceptible | Resistant | Test set pool |
| --- | --- | --- | --- | --- | --- |
| Rifampin | *rpoB* | 3102 | 2792 (90%) | 310 (10%) | 500 (400 susceptible, 100 susceptible) |
| Isoniazid | *katG* | 3138 | 2559 (81.5%) | 579 (18.5%) | 500 (400 susceptible, 100 susceptible) |
| Pyrazinamide | *pncA* | 3084 | 2938 (95.3%) | 146 (4.7%) | 500 (400 susceptible, 100 susceptible) |

Until the recent publication of the large-scale study by Walker *et al.* (the Global Dataset in our research), *M. tb* drug resistance associated studies usually had a sample size of a few to a few hundred strains [10]. In addition, there are not such large-scale studies available for other infectious agents yet. Thus, it is necessary to investigate how Composite MetaDistance would perform with isolated outbreaks. An Outbreak Dataset of 161 strains consisting of strains from an outbreak in China was curated [30]. Drug resistance to Rifampin was predicted using a per-gene per-drug feature set for *rpoB*. In contrast to the Global Dataset, the Outbreak Dataset was highly imbalanced with 117 strains resistant and 44 strains susceptible to Rifampin. Resistance to Rifampin was predicted using a feature set for its drug target *rpoB*. The training set consisted of 141 strains with a balanced test set of 20 strains set aside for evaluation.

### Mutation detection pipeline

Mutation identification and analysis used a modification of a computational pipeline previously introduced in [33]. Fastq files for *M. tb* strain sequences were mapped to the H37Rv reference genome using BWA-MEM [34]. The H37Rv genome was downloaded from NCBI [35]. Strains with multiple fastq files (and thus multiple SAM alignments from BWA-MEM) were combined using Picard-Tools [36]. Single nucleotide variants (SNV) were identified with samtools 0.1.18 and bcftools 0.1.17-dev [37, 38]. Only SNVs with a minimum phred score of 20 and a minimum 20x high-quality read depth were used for downstream analysis and prediction. A further filtering step was applied to include only SNVs with the ratio of high quality reads to all reads > 0.6, and at least 3x more alternate calls than reference calls. The filtered SNVs were then functionally annotated using ANNOVAR with the H37Rv reference genome, and used for feature creation [39]. The pipeline is available at github (https://github.com/dattagargi/MutationAnalysisPipeline), and a version of this integrated bioinformatics pipeline has been previously published [33].

### Comparing Composite MetaDistance with existing algorithms

We used both the Outbreak Dataset (161 strains) and the Global Dataset (3650 strains) to compare Composite MetaDistance to MetaDistance and existing classical machine learning methods like random forests, SVM with a radial basis function (rbf) kernel and KNN. For MetaDistance, we used *p*=1. We used the the scikit-learn 0.16.1 implementation of random forests, SVM and KNN with default parameters. To calculate the best parameters for the SVM rbf kernel, we performed a grid search of possible values for gamma (10^−9^ to 10^3^) and C (10^−2^ to 10^10^), and evaluated on the training set with 5-fold cross validation. Composite MetaDistance was also compared to the modified KNN (mod-KNN) algorithm for unbalanced datasets used in MetaDistance.

#### Random subsampling

The Global Dataset is a recently published dataset with 3,650 strains [10]. Until this dataset, it was di cult to obtain genomic information for more than a few hundred strains of *M. tb*. It is similarly difficult to obtain genomic data for other infectious agents as well. In addition, for second-line drugs, even the Global Dataset has phenotypic information for only a few hundred strains. Thus, it is important to evaluate how Composite MetaDistance scales down. To evaluate the robustness of Composite MetaDistance to training set size, the 3,650-strain Global Dataset was randomly subsampled to create training sets of 20, 50, 100, 500 and 1000 samples. This was done 20 times to evaluate Composite MetaDistance.

### Evaluation

The training and validation set splits for each per-gene per-drug resistance prediction problem, and the evaluation metrics are detailed in this subsection.

#### Training and test sets

For each drug tested, only a subset of the 3,650 strains in the Global Dataset had phenotypic susceptibility information available. 500 randomly selected strains were set aside to create test sets (400 susceptible, 100 resistant). From this pool of 500 strains, 20 balanced test sets of 100 strains each (50 susceptible, 50 resistant) were created with random subsampling. All algorithms and training set subsamples were tested on these 20 test sets to evaluate their performance. The rest of the strains with phenotypic susceptibility for each of the first-line drugs were used as the training set strains for each gene for the drug, and for the combined resistance profile wherever applicable (Table 2).

#### Evaluation metrics

To evaluate the performance of Composite MetaDistance, and compare it against existing methods, we used prediction error for model performance. The prediction error of a model is (1- accuracy of the model). Thus, lower prediction error means higher accuracy [24].

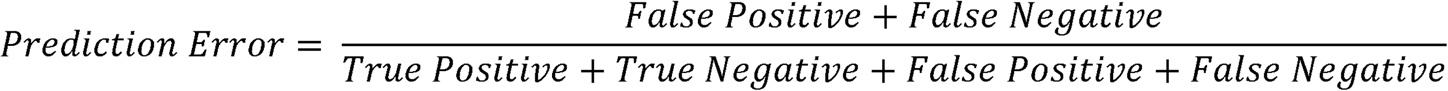

For the global dataset, the final evaluation metric was the median test set prediction error over the 20 random test sets. When Composite MetaDistance was trained on subsamples of the global dataset, the final evaluation metric was the median test set prediction error over the 20 learned instances for a given sample size, each tested on the 20 random test sets.

We also calculated the sensitivity, specificity and F1 score of each algorithm. As with the prediction error, for the global dataset, the final reported metric was the median test set prediction error over 20 random test sets. Metrics for the random subsampling cases were the median test set prediction error of all 20 trained learners for a given sample size, each tested on the 20 test sets.

## Results

Composite MetaDistance, the algorithm developed to predict per-gene per-drug resistance, was compared against existing machine learning algorithms to evaluate its robustness and accuracy. Prediction error was used as the primary metric for these evaluations, with sensitivity, specificity and F1 scores reported. The Global Dataset consisted of 3,650 publicly available strains of *M. tb*. Three prediction tasks were performed with this dataset: 1. Predict resistance to Rifampin due to mutations in the gene *rpoB*, 2. Predict resistance to Isoniazid due to mutations in *katG*, and 3. Predict resistance to Pyrazinamide due to mutations in *pncA*. In addition, drug resistance to Rifampin was first predicted using only strains from the Outbreak Dataset, without any knowledge of the Global Dataset. The Outbreak Dataset represents a real-world outbreak scenario, where not a lot of information is easily available to model drug-resistance.

### Outbreak dataset: 161 strains

In the outbreak dataset, we had a small sample classification problem, with only 141 *M. tb* strains to use as our training set. Composite MetaDistance was evaluated against existing machine learning methods, with a balanced test set of 20 strains (10 resistant, 10 susceptible). The classification task was to predict whether a strain of *M. tb* was resistant or susceptible to Rifampin, based on features in the gene *rpoB*. We first constructed a feature set for the prediction task, based on mutations in *rpoB* for the strains. This multi-view feature set included 1172 amino acid-based, 1172 structure-based, 6 function-based, 1 similarity-based and 1 phylogeny-based features. The features were of mixed attribute types, with 1173 continuous, 1172 ordinal and 7 nominal features (Table 1). Composite MetaDistance outperformed other methods, with a median prediction error of 0.3 and F1 score of 0.667 (Fig. 4A). The base algorithm, MetaDistance on the other hand had an error of 0.4. Since the original application of MetaDistance to metagenomics used proportion data scaled between 0.0 and 1.0, we tested whether MetaDistance applied to unscaled feature data in this context was unfairly compared against Composite MetaDistance, which uses range-normalized features. However, applying MetaDistance to range-normalized features did not improve its performance, thus highlighting the superior prediction accuracy of Composite MetaDistance.

**Figure 4.**
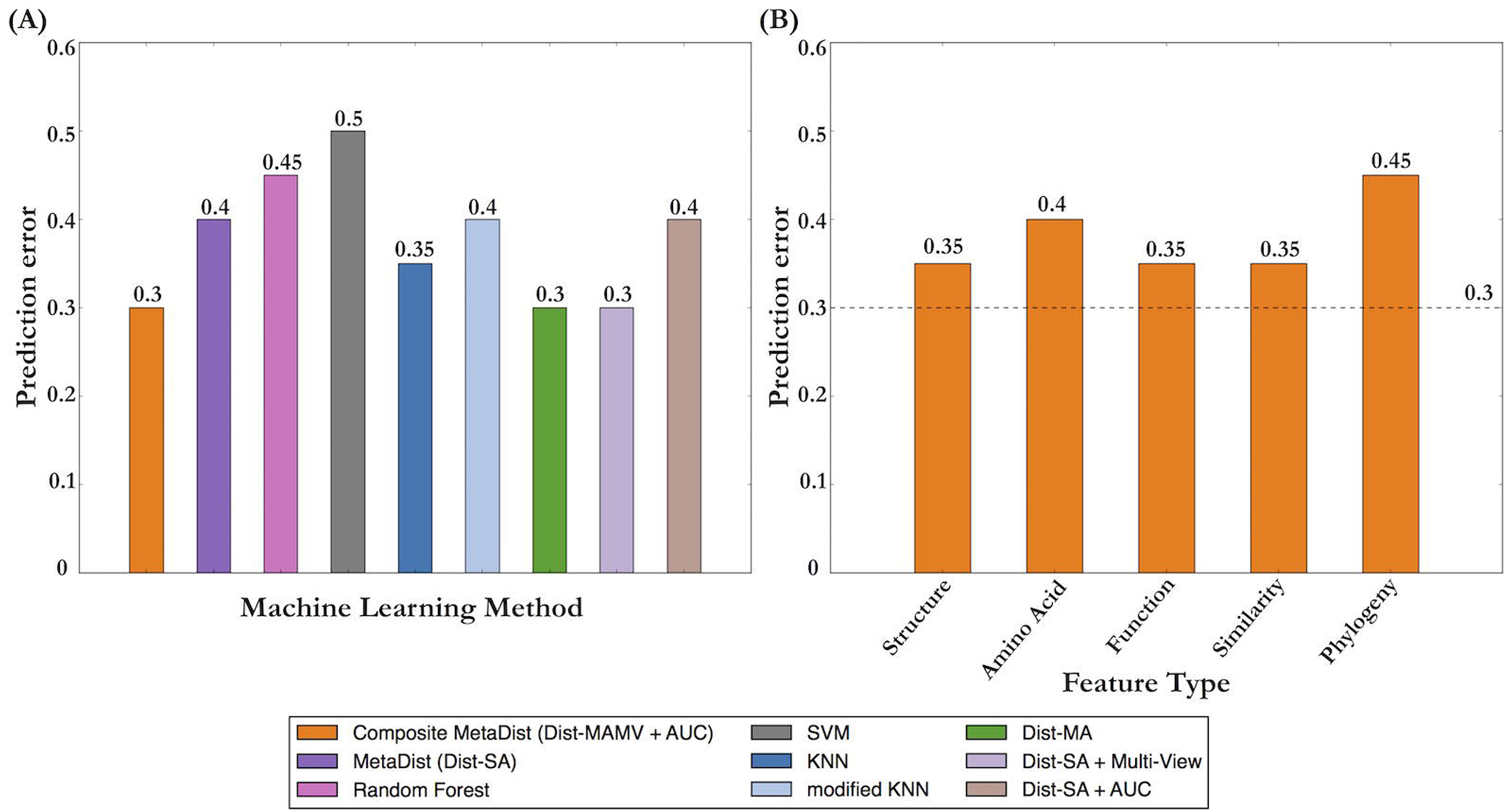
Outbreak dataset results. The outbreak dataset consisted of 141 training samples and 20 test samples. A. Comparison of machine learning methods on the outbreak dataset. Composite MetaDistance (in orange) outperforms all other methods. B. Individual feature view analysis with Composite MetaDistance on the outbreak dataset. No single view performs as well as the combination of all views.

Specificity, sensitivity and F1 score analyses for each algorithm for the Outbreak Dataset were performed (Table 3). While Composite MetaDistance, MetaDistance and modified KNN had a lower sensitivity (0.4) than other algorithms, Composite MetaDistance had no false positives, thus resulted in a specificity of 1.0 and correctly identified all negative instances. Random Forest, SVM and KNN had very high sensitivity and very low specificity scores, thus indicating a high bias towards false positives. This is not desirable, since they would incorrectly identify (almost) all examples as resistant to the first line drug Rifampin, thus resulting in the unnecessary prescription of the more toxic second line drugs to individuals with Rifampin-susceptible cases of TB [5]. This might in turn also result in an increase in potential for future acquired resistance [5].

**Table 3.**
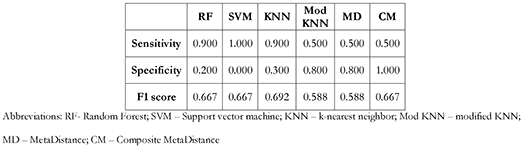
Sensitivity, Specificity and F1 scores for the Outbreak dataset

#### Composite MetaDistance component analysis

To investigate which component of Composite MetaDistance contributed a performance boost, we evaluated each component of Composite MetaDistance (*dist*_*MA*_; multi-view step; and pre-initialized weights from individual AUC analysis) individually with the base MetaDistance algorithm (Fig. 4A). Adding only a mixed attribute distance function (*dist*_*MA*_) to MetaDistance showed a drop in prediction error, from 0.4 for MetaDistance to 0.3 with MetaDistance + *dist*_*MA*_, thus demonstrating the power of proper handling of mixed attribute features. Even with only a few nominal features (0.3%, Table 1), proper handling improved the prediction tremendously, especially for a small training set of 141 strains. Adding a multi-view component to the base algorithm also showed a drop in prediction error (from 0.4 in MetaDistance to 0.3 with multi-view learning), while adding pre-initialized feature weights to MetaDistance did not improve performance (Fig. 4A). The improvement in performance by adding the multi-view component demonstrates the benefits of removing bias for larger feature views. Though not hugely impacting the prediction error, adding pre-initialized feature weights showed a speed up over the base algorithms convergence rate from 53 iterations in MetaDistance to 28 with pre-initialized weights.

#### Individual view analysis

We performed a view-level analysis to determine whether prediction with one view performs as well as the combination of all views with Composite MetaDistance. Our results showed that Composite MetaDistance with all views (dashed line in 4B), and thus all available information encoded, outperformed all individual views (Fig 4). The results of our individual view analysis demonstrate that data from a combination of sources makes more accurate and robust predictions. Moreover, the views contributing few features, such as the 1-feature similarity-based view and the 6-feature function-based view, prove to be extremely informative, especially in comparison to the structure or amino acid-based views with >1000 features each. This motivates our use of multi-view coefficients to balance information from all feature views.

### Global dataset: 3650 strains

The global dataset consisted of 3650 publicly available strains of *M. tb*, described in the methods. Three prediction tasks were performed with this dataset: 1. Predict resistance to Rifampin due to mutations in the gene *rpoB*, 2. Predict resistance to Isoniazid due to mutations in *katG*, and 3. Predict resistance to Pyrazinamide due to mutations in *pncA*.

#### Rifampin

Rifampin binds to the β-subunit of RNA polymerase in *M. tb*, and mutations in the gene *rpoB* are primarily responsible for resistance to the drug [5, 27, 31, 53]. For *rpoB*, the binding site for Rifampin is a well-defined 81bp region between amino acids 507-533, the RRDR [40]. Mutations in the 81bp Rifampin-resistance determining region (RRDR) of *rpoB* account for over 95% of Rifampin-resistant strains of *M. tb* [27, 31, 53]. The classification task was to predict whether a strain of *M. tb* was resistant or susceptible to Rifampin, based on features in the gene *rpoB*. A feature set was first constructed, based on mutations in *rpoB* for strains with phenotypic susceptibility information available for Rifampin. The structure features were calculated as the Euclidean distance of mutations from the computed center of the binding site. Since *rpoB* has 1172 amino acids, the multi-view feature set included 1172 amino acid-based, 1172 structure-based, 6 function-based, 1 phylogeny-based and 1 similarity-based features (Fig. 3). The features were of mixed attribute types, with 1173 continuous, 1172 ordinal and 7 nominal features (Table 1). The training set consisted of 3102 strains (Table 2).

#### Isoniazid

Isoniazid requires activation by the catalase/peroxidase enzyme KatG, which is encoded by the gene katG [53]. The mutation S315T in *katG* accounts for 40-64% of Isoniazid-resistant strains in the general population. The main Isoniazid binding site in *katG* is in the heme-binding channel of the protein between amino acids 104 and 381, and structure features were calculated as the Euclidean distance from the center of this binding site [41, 42]. The feature set for *katG* for the per-gene per-drug prediction included 740 amino acid-based, 740 structure-based, 3 function-based, 1 phylogeny-based and 1 similarity-based features. Thus, there were 741 continuous, 740 ordinal and 4 nominal features (Table 1). The training set consisted of 3138 strains (Table 2).

#### Pyrazinamide

Pyrazinamide needs to be converted to pyrazionic acid by the enzyme pyrazinamidase (PZase), which is encoded by the gene *pncA* [53]. Loss of PZase activity is associated with Pyrazinamide resistance, and mutations in *pncA* have been found in clinically resistant strains [29, 53]. The Pyrazinamide binding site in *pncA* is a small crevice 10Å by 7Å between amino acids 8 and 138 [29]. The *pncA* feature set included 186 amino acid-based features, 186 structure-based features, 1 function-based feature, 1 phylogeny-based feature and 1 similarity-based feature. The training set had 3084 strains.

#### Prediction results

As with the outbreak dataset, we evaluated Composite MetaDistance against other machine learning methods. Each algorithm was trained for a given prediction task on an unbalanced training set of ~3000 strains classified as either resistant or susceptible to the respective drug (Table 2). Each algorithm was tested on 20 balanced test sets of 100 strains each (50 resistant, 50 susceptible) (Table 2). Median prediction error across the 20 test sets for Rifampin, Isoniazid and Pyrazinamide resistance prediction are shown in Fig. 5A, B and C respectively. Composite MetaDistance (shown in orange) consistently performed as well or better than all other algorithms across all three prediction tasks.

**Figure 5.**
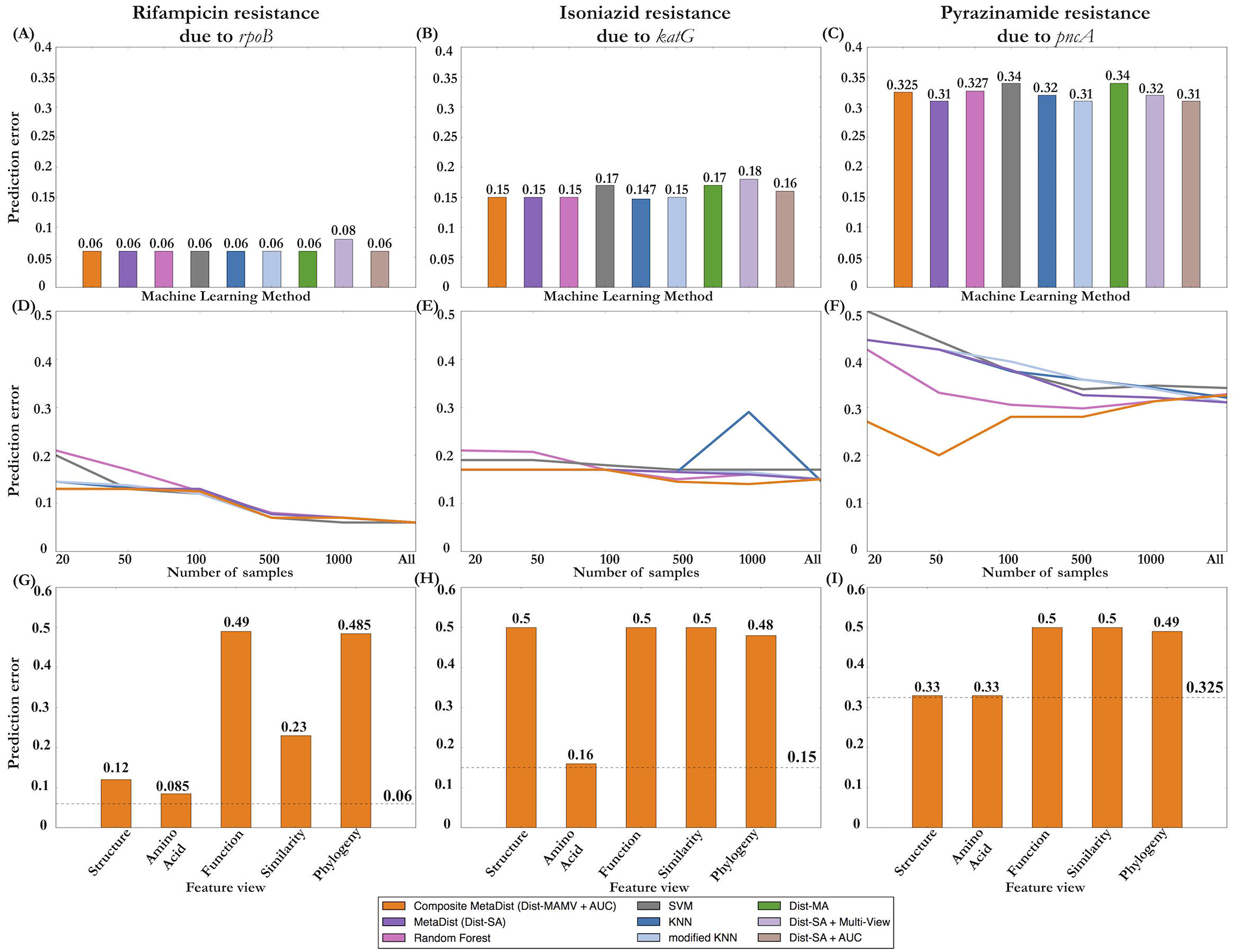
Global dataset. Results for three prediction tasks (resistance to Rifampin, Isoniazid and Pyrazinamide) for the global dataset. The left panel (A, D, G) shows results for resistance predictions to Rifampin. Middle panel (B, E, H) shows results for resistance prediction to Isoniazid. Right panel (C, F, I) shows results for resistance prediction to Pyrazinamide. Top row (A, B, C) shows prediction results for the whole training set, and comparisons to other algorithms. Middle row (D, E, F) shows subsampling results for each prediction task. Bottom row (G, H, I) depict individual feature view analysis results for each prediction task.

In addition to the whole Global Dataset, random subsampling was done within the aforementioned prediction problems to evaluate the robustness of Composite MetaDistance for small sample prediction problems. Specificity, sensitivity and F1 scores of all algorithms on all ~3000 strains are given in Table 4 (bottom row each section). Specificity across most prediction tasks and machine learning algorithms was 1.0. Thus, the algorithms were very good at predicting negative (susceptible) examples. Sensitivity and F1 score varied, and followed the trends of prediction error where Composite MetaDistance did as well as or better than the other algorithms when trained on all ~3000 strains. We performed Mann Whitney U tests to compare Composite MetaDistance to existing algorithms, and the p-values are reported in Supp. Table 1.

**Table 4.**
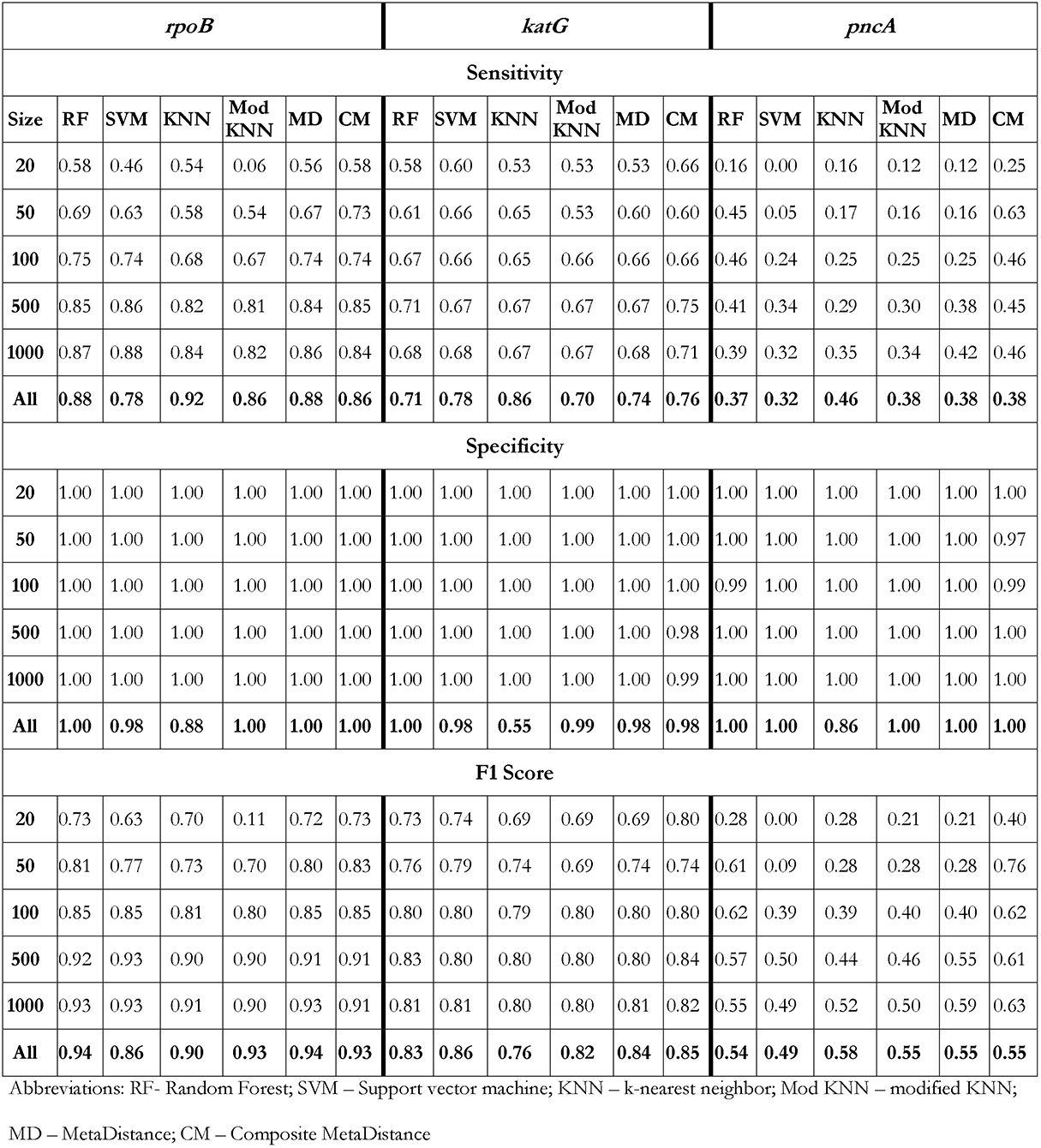
Sensitivity, Specificity and F1 Scores for the Global dataset

#### Composite MetaDistance component analysis

We evaluated the performance of individual components of Composite MetaDistance added to the base MetaDistance algorithm, for the entire training set, for each prediction task (Figs 5A, 5B, 5C). For predicting drug resistance to Rifampin due to mutations in *rpoB*, although there was no significant improvement in prediction error upon adding the individual components, there was an improvement in computational speed. Computational speed here was determined by the number of iterations of the SVM conjugate gradient training needed for convergence to the final learned weights. Adding *dist*_*MA*_ to MetaDistance reduced the number of iterations taken by 50%. For predicting resistance to Isoniazid due to mutations in *katG*, adding a mixed attribute distance metric reduced the number of iterations taken to 34% of the number of iterations for MetaDistance, and adding a multi-view step sped up the base MetaDistance algorithm by reducing iterations by 67%. adding individual AUCs as pre-initialized weights reduced the number of iterations by 49% of the number of iterations for MetaDistance. Adding *dist*_*MA*_) did not speed up the algorithm by a significant amount. These improvements in speed occurred as a result of faster convergence by the conjugate gradient descent in MetaDistance.

#### Sample size analysis

To evaluate whether the algorithm is generally robust for small sample sets, the Global Dataset was randomly subsampled for 20, 50, 100, 500 and 1000 sample size sets, and trained on these smaller sample sets. Random subsampling with replacement was done 20 times for each sample size, and each model was tested on the same 20 test sets used in Figs 5A-5C. The median prediction error for each sample size, including the whole training set of size ~3000 for each prediction task is plotted in Figs 5D, E and F, for each machine learning algorithm tested. The complete error distributions for Rifampin, Isoniazid and Pyrazinamide prediction tasks for all sample sizes are depicted in Supp. Fig. 1, 2 and 3 respectively. Composite MetaDistance (shown in orange) consistently performed as well or better than all other algorithms for both small and large training sets (Fig 5D, 5E, 5F). For sample size of 1000 or less, Composite MetaDistance had the best performance, thus further demonstrating the robustness of the algorithm (Fig 5D, 5E, 5F). Furthermore, the curves plotted in Figures 5D,E,F are flattest for Composite MetaDistance. Thus, Composite MetaDistance is sample size agnostic and can be used for small sample machine learning problems, such as predicting drug resistance to the second-line drugs or drug resistance prediction in other infectious agents such as NTM. Thus, Composite MetaDistance is sample size agnostic and can be used for small sample machine learning problems.

As with the Outbreak Dataset, in order to confirm that we were not penalizing the base algorithm, MetaDistance, for its use on non-scaled features, we applied MetaDistance to range-normalized features in one of the prediction tasks (pyrazinamide prediction due to mutations in *pncA*). MetaDistance using range-normalized features performed worse than MetaDistance or Composite MetaDistance across almost all training sample sizes (except 100). The resulting median prediction errors are shown in Supp. Fig. 4.

Specificity, sensitivity and F1 scores of Composite MetaDistance compared to other algorithms for increasingly larger subsampled training sets are given in Table 4. As with the models trained on all ~3000 strains, specificity for subsampling was 1.0 or nearly 1.0 for almost all training sets, algorithms, and training set sizes. Thus, predicting resistance is a more difficult task than predicting susceptibility for the global dataset, as expected given the imbalance of a few resistant strains. Sensitivity and F1 scores follow the trends of prediction error.

#### Individual view analysis

A view-level analysis was performed to determine whether prediction with one view performs as well as the combination of all views with Composite MetaDistance. For the Global Dataset, an individual view analysis was performed for each of the three prediction tasks, as shown in Fig. 5G, H and I. No single view outperformed Composite MetaDistance (dashed horizontal line). In contrast to the outbreak dataset, the structure and amino acid-based features prove to be among the most informative for the global dataset. These results, and the fact that no one feature performs as well as Composite MetaDistance, further demonstrated that data from a combination of sources makes more accurate and robust predictions.

#### Difference in prediction accuracy across the three tasks

The overall prediction error is much lower for Rifampin susceptibility prediction than Isoniazid or Pyrazinamide. Similarly, sensitivity and F1 scores are generally higher for Rifampin prediction than Isoniazid or Pyrazinamide. This is likely due to the fact that a significant amount is known about how and where Rifampin binds to *rpoB*. *rpoB* is a RNA polymerase protein with a well-defined binding site [48]. In general, the farther away a mutation is from the binding site, the lower the mutation’s effect on drug resistance. Around 95% of strains resistant to Rifampin in nature have mutations in the RRDR of *rpoB*, which is captured in the weights learned by Composite MetaDistance. Mutations in the RRDR of *rpoB* in the Global Dataset are very good predictors of resistance.

On the other hand, Isoniazid resistance-conferring mutations can occur throughout the *katG* protein, likely disrupting a heme-binding channel involved in the activation of Isoniazid [41, 42]. Since the protein structure is not circular like *rpoB*, and the mode of action and mechanisms of resistance are complex and incompletely understood, accuracy of Composite MetaDistance for this prediction task was lower than the *rpoB* prediction task. Pyrazinamide resistance-conferring mutations can occur throughout *pncA*, in part since it is a small protein (186 aa), and mutations in positions far away from the binding site can disrupt the configuration thought to be involved in the activation of Pyrazinamide [29]. Thus, the prediction accuracy is worse than Rifampin or Isoniazid.

Composite MetaDistance outperforms other algorithms by a non-trivial margin for Pyrazinamide resistance prediction, demonstrating the efficacy of our algorithm with noisy data and small sample sets.

## Discussion

The paper introduces a novel method to predict drug-resistance in *M. tb*, with a more robust and innovative feature set than devised before. The primary dataset used for these predictions is a large-scale study of 3,650 strains of *M. tb* from many countries [10]. This was the largest study for *M. tb* to date and provided a rich database for our research. For drug resistance prediction, we first motivated the need for a novel feature set to characterize mutations and incorporate additional information. Such a robust and comprehensive feature set has not been defined and used in drug resistance prediction prior to this research. The next step in the process is to build a machine learning model that can properly handle such complex datasets and accurately predict drug resistance. To this end, we have created Composite MetaDistance, a simultaneous feature selection and classification algorithm for mixed, multi-view unbalanced genomic data. Our algorithm deals with many challenging data issues in machine learning. The feature set per gene contains mixed attribute features from multiple data sources (feature types or views), on the order of a few hundred to a few thousand features per gene. Drug resistance data is also usually extremely skewed, with 90% of strains in the real world being susceptible to a particular drug. We have devised a composite distance function, called the Mixed Attribute Multi-view distance (*dist*_*MAMV*_) to properly deal with such datasets. By combining *dist*_*MAMV*_ with MetaDistance, a method in metagenomics for classification of high dimensional unbalanced datasets, we created Composite MetaDistance, a sample size agnostic and robust prediction algorithm to classify mixed attribute data from multiple data sources, especially nominal features [24]. Composite MetaDistance outperforms existing algorithms for small sample training sets. This is important for resistance prediction for drugs without a lot of training samples, and for other infectious agents with a dearth of data available. It generally outperforms existing methods or performs as well as the best method even with less than 1% nominal features. We expect Composite MetaDistance to perform even better with a higher percentage of nominal features.

### Robust feature set

Until now, drug resistance prediction tasks for most diseases use feature sets made of either sequence- or structure-based features for each target gene. The feature set presented in this research is rich and diverse, and incorporates different types of features (feature views). The amino acid sequence-based features convey the likelihood of an amino acid change in nature (BLOSUM62 values), thus effectively encoding the impact of a mutation on the protein. The structure-based features encode information about the spatial distance of a mutation from the binding site. Function-based features depict whether a functional domain in the protein is modified, thus having implications on the function of the protein. The similarity-based feature is the divergence of the gene sequence from the susceptible H37Rv reference genome. Strains with a higher divergence from H37Rv are more likely to be resistant. The phylogeny-based feature calculates the closest reference strain for the strain in question. Some reference strains are more virulent than others, and some have a higher probability of gaining resistance than other reference strains. This information is encoded in the phylogeny-based feature. Thus, the feature set not only derives novel feature views from sequence mutations, but also encodes potentially useful prior and publicly available information, thus making it more robust and informative than existing feature sets in *M. tb* research (Fig. 3). Moreover, the combination of feature views was found to be a better predictor than any single feature type (Figs. 4, 5). Such a feature set has not been devised for drug resistance prediction before, and has the potential to improve drug resistance prediction immensely. It can also be easily extended to include even more information, such as binding affinities. Depending on the gene, this feature set contains anywhere from a few hundred to a couple thousand features, with features from different views (feature types) and different data types (continuous, ordinal, nominal).

The sequence- and structure-based views performed worse in the smaller Outbreak Dataset than the Global Dataset (Tables 3,4 and Fig. 4,5). The difference may be attributable to the fact that for these two large feature views, the true signal could be better captured in the larger Global training set than in the smaller Outbreak Dataset, whereas for the Outbreak Dataset, the similarity and phylogeny views with a few features allowed a succinct summarization of the limited training data. These results highlight the importance of incorporating multiple views in the feature set.

### Small-sample classification

Composite MetaDistance consistently outperformed existing machine learning algorithms with sample sizes less than 1000 in the Global Dataset, and in the Outbreak Dataset, which is a small-sample classification problem. In the Global Dataset, while specificity was around 1 for all algorithms, Composite MetaDistance was more sensitive to detecting resistant strains than other algorithms for small sample sizes (Table 4). In addition, as shown in Fig. 4, the curves for Composite MetaDistance were flattest across sample sizes for each prediction problem, thus demonstrating the sample-size agnostic behavior of our algorithm. Both the Global Dataset random subsampling and Outbreak Dataset analysis demonstrate the power of Composite MetaDistance over existing algorithms for small sample machine learning problems. This is very important for classifying resistance in isolated, and for other infectious diseases, as specified in further applications.

### Mixed attribute distance function

As seen in our datasets, most real world applications of machine learning deal with features of mixed attribute types. Composite MetaDistance accounts for differences in mixed data by defining a mixed-attribute distance metric, which is an extension of the Gower similarity index. The distance metric used in the base algorithm (*dist*_*SA*_) works well for continuous or ordinal data, but will give nonsensical distances for nominal features [14, 15]. The mixed attribute distance (*dist*_*MA*_) deals with this issue by defining a binary distance metric for nominal features. Our feature sets are high-dimensional with thousands of features. For each gene (*katG*, *pncA*, *rpoB*), the number of features of different data types is shown in Table 1. Even with a much higher number of continuous and ordinal features than nominal features, several of which are binary nominal features, Composite MetaDistance outperforms other algorithms. This demonstrates the power of proper handling of mixed attribute types. For datasets with a higher dimensionality of nominal datatype features, especially those with a cardinality of more than two, we expect our algorithm to outperform existing algorithms to an even greater extent.

### Multi-view coefficients and pre-initialized feature weights

In Composite MetaDistance, the distance function is calculated as a linear combination of distances per feature (*dist*_*MA*_). The multi-view learning step further normalizes the distances, so that each view has an equal contribution to the overall distance function, regardless of the number of features in the view. This is done by a weighted linear combination of individual feature-view distances, thus coming up with a mixed attribute multi-view function (*dist*_*MAMV*_). The view coefficients are currently calculated as an inverse of the cardinality of the view, which ensures that a view with a large number of features (for example, structure features in our data) does not overwhelm a view with a small number of features (phylogenetic features for example). The benefits of removing the bias for larger feature views is seen in Fig 4. A in the context of MetaDistance, where adding just the multi-view component improved MetaDistance performance. The multi-view component in conjunction with the mixed attribute component of the composite distance function gives Composite MetaDistance the boost it needs to outperform other algorithms. Pre-initializing feature weights as normalized individual AUCs from a random forest analysis informs initial weights per feature. These weights (*w*_*m*_) are initialized as 1.0 in MetaDistance and are learned using a conjugate gradient descent algorithm with the quadratic SVM [24]. Having informed initial weights instead of 1.0 decreases the number of iterations needed for convergence of the conjugate gradient descent, and thus reduces the amount of time needed to learn the feature weights for the final classification task. This speeds up the algorithm by a considerable amount. Both the multi-view coefficients and pre-initialized feature weights can be modified to fit different feature sets and feature types. For example, in a dataset where there is not a large discrepancy between feature view sizes, or where one feature from a large view is more important than all other features, one can envision adjusting the view coefficients to reflect prior information. These modified view coefficients would then be used with Composite MetaDistance to predict samples.

### Modified k-nearest neighbor (KNN) algorithm

A regular KNN predicts the class of a sample by polling the closest *k* neighbors and assigning the class with the maximum number of samples in the neighborhood [49]. Although this works for balanced datasets, as datasets become more and more unbalanced, a regular KNN starts misclassifying samples as the larger class, even when using weighted distances [50]. For example, in the global dataset where >80% of the samples are susceptible to any given drug, any neighborhood of k samples will have a majority labeled susceptible. Thus, a regular KNN would misclassify any resistant sample. To deal with the problem of class imbalance, MetaDistance uses a modified KNN algorithm. The modified KNN calculates the *k* closest neighbors to a sample in each class, and classifies the sample in the class with the closest mean neighborhood score, thus dealing with imbalance in data. This results in higher prediction accuracy as compared to a regular KNN, as demonstrated in Fig. 5 (KNN in dark blue, modified KNN in light blue).

### Further applications

Composite MetaDistance is a robust algorithm for classification of high dimensional unbalanced data with mixed-attribute multi-view features, which is the case for many real-world datasets. Thus, Composite MetaDistance can be used to classify any dataset that has similar properties and is not limited to genomic or biological data. An advantage of our per-gene per-drug resistance prediction and combined resistance prediction pipelines over existing molecular assays and genomic resistance identification methods is that they are not restricted to finding particular mutations in a gene, but take into consideration all possible mutations in candidate genes. This is especially important for cases where exact drug resistance conferring mutations may not be known for the gene. For example, for Bedaquiline, Clofazimine and Cycloserine, some of the target genes in *M. tb* are known, but exact mutations in target genes that confer resistance are largely unknown [52, 53]. Current molecular assays, genotypic and sequencing based methods would thus not be able to identify resistance to these drugs, but Composite MetaDistance would be able to identify patterns associated with resistance, and predict resistance. The per-gene feature set for *M. tb* is also applicable for any infectious disease.

Composite Distance may also be used for feature selection, as the weights from the SVM can be used to find important features for the classification task at hand. For example, for Isoniazid prediction using *katG*, in addition to S315T, the weights learned by Composite MetaDistance highlighted a few additional amino-acid positions with a high correlation to resistance. The amino acid position with the highest feature weight was position 35, followed by positions 241 and 203. Similarly, drug-resistance prediction to Pyrazinamide highlighted amino acid positions 71, 133 and 96. These positions are around the binding site for Pyrazinamide and it is important to investigate them in the lab for further implications to resistance. Further investigation into the features (and thus amino acid positions) highly weighed by Composite MetaDistance could shed light on individual mechanisms for resistance, which can then be confirmed in the lab.

An alternative approach to predicting drug resistance for relatively well-studied drugs such as Rifampin would be to create a hybrid classifier with an initial decision tree to detect known mutations for resistance, such as mutations in the RRDR. This could be the simple molecular classifiers for those drugs. If a strain has a mutation in the region, it is classified as resistant. Otherwise, the machine learning pipeline is used to classify the strain as resistant or susceptible. An analysis was performed in the Global Dataset to predict drug resistance using only features (positions and mutations) outside the Rifampin-resistance determining region (RRDR). The median prediction error when RRDR was not considered to predict drug resistance was still 0.06, thus indicating that Composite MetaDistance is a good predictor of drug resistance in cases where a molecular method might be inefficient in detecting drug resistance. Composite MetaDistance is extensible to any drug resistance prediction task and as shown in our results works especially well with small training samples (Fig 4, Fig 5D, E, F). This is very important for resistance prediction with second and third line drugs, which usually have a smaller number of available resistant samples, and other diseases in which data is sparse. Many diseases (such as NTMs) do not have many publicly available sequenced strains, yielding small training sets. Using traditional algorithms for resistance prediction in these cases may cause noise and overfitting [24]. Composite MetaDistance on the other hand is robust for training sets as small as 20 to 50 samples, and thus can be used for small-sample classification. Furthermore, Composite Distance can be used for any multi-label classification task for high-dimensional mixed data, with or without multiple feature types and redundant features.

In contrast to the Global Dataset, the Outbreak Dataset is an anomalous dataset, with around only 33% of the resistant strains having a mutation in the Rifampin-resistance determining region, and serves an indicator of the diversity of outbreaks around the world. The fact that Composite MetaDistance outperforms other algorithms in the Outbreak Dataset demonstrates that Composite MetaDistance can be used to predict drug resistance for isolated outbreaks of infectious agents.

### Limitations

The primary limitation of Composite MetaDistance is that due to the severe penalizing of view-coefficients of large feature views, features in those views that have implications for resistance may be unfairly suppressed. This issue can be circumvented by devising view coefficients (either learned or static) that have lower penalties, inflating individual feature coefficients for the feature(s) in question to counter the penalty, or creating a new feature view with just the features that are known to be important for the predictions. The view coefficients can be learned based on the feature set and used for Composite MetaDistance. Another possible limitation is that using the view coefficients for predicting test samples after the feature weights have been learned by conjugate gradient training may in effect nullify the effectiveness of the learned weights. Thus, an important future experiment is to create predictions using only the learned feature weights. These limitations also highlight the strength of Composite MetaDistance in that once a modified view coefficient is defined, it can be plugged into the composite distance function without changing the algorithm. Thus, the individual components are modular and customizable.

## Conclusions

The first step in a machine learning prediction problem is to create a robust and informative feature set. The feature set specification presented in this paper utilizes mutational information and publicly available information for each gene, and is much richer than ever devised previously for drug-resistance. The next step in the process is to build a machine learning model that can properly handle such complex datasets and accurately predict per-gene per-drug resistance. To this end, we have created Composite MetaDistance, a simultaneous feature selection and classification algorithm for mixed, multi-view unbalanced genomic data. Composite MetaDistance is sample size agnostic and has high accuracy for smaller training sets, thus making it possible to explore datasets with a scarcity of information available. For example, when detecting novel mechanisms for drug resistance, especially for newer drugs, the training set will be small since not a lot of phenotypic information would be known. Composite MetaDistance can be used in these cases to accurately predict drug resistance. In addition, an exploration of the feature weights learned by Composite MetaDistance can identify important mutations for drug resistance, thus indicating novel mechanisms that can be investigated further in a wet lab. Thus, Composite MetaDistance makes robust predictions with small training samples, and can be used to make inferences for novel mechanisms of drug resistance with small training sets.

## List of abbreviations

*M. tb*: *Mycobacterium tuberculosis*
TB: Tuberculosis
ML: Machine Learning
WHO: World Health Organization
WGS: Whole Genome Sequencing
RF: Random Forest
KNN: K-Nearest Neighbor
SVM: Support Vector Machine
AUC: Area Under the receiver operating characteristic Curve
SNV: Single Nucleotide Variant
rbf: Radial Basis Function
MD: MetaDistance
CM: Composite MetaDistance

## Declarations

### Ethics approval and consent to participate

Not applicable

### Consent for publication

Not applicable

### Availability of data and material

The datasets used for this research are publicly available, and enumerated in Walker *et al.* and

### Authors’ contributions

GD and SL wrote the main manuscript, with additional ideas from MS. GD, SL and MS conceived the project. GD and SL developed methods and designed experiments. GD and MS conceived and designed the feature vectors. GD analyzed all strains and created the feature vectors. NH created phylogeny-based features. GD implemented and validated the methods, and ran experiments. GD, SL and MS interpreted results. GD generated figures and tables for manuscript. All authors reviewed and approved the final manuscript.

## Acknowledgements

We would like to thank Dr. Lawrence Hunter for his advice with the methods, and Benjamin Garcia for his help with similarity-based feature creation. We would also like to thank the Computational Bioscience Program at the University of Colorado School of Medicine, and National Jewish Health for the resources to conduct this study.

