## Supplemental Information for "A composite method to infer drug resistance with mixed genomic data"

**Supp. Table 1. Mann-Whitney U test p-values for Composite MetaDistance vs. existing algorithms**

| **Drug** | **# samples** | **MetaDist** | **RF** | **SVM** | **KNN** | **Mod KNN** |
| --- | --- | --- | --- | --- | --- | --- |
| **Rifampin** | 20 | 0.33 | 0.22 | 0.13 | 0.12 | 0.18 |
|  | 50 | 0.25 | 0.47 | 0.18 | **0.02** | **0.02** |
|  | 100 | 0.24 | 0.42 | 0.09 | **0.0009** | **0.0007** |
|  | 500 | 0.1 | 0.5 | 0.02 | **0.0002** | **0.0016** |
|  | 1000 | 0.38 | **0.03** | 0.07 | 0.1 | **0.0001** |
|  | All | 0.1 | 0.17 | **<0.0001** | **<0.0001** | 0.39 |
| **Isoniazid** | 20 | 0.26 | 0.45 | 0.07 | 0.04 | 0.1 |
|  | 50 | 0.42 | 0.25 | 0.12 | **0.02** | **0.006** |
|  | 100 | 0.42 | 0.28 | 0.35 | **0.002** | 0.17 |
|  | 500 | **0.001** | 0.11 | **0.0004** | **0.001** | **0.001** |
|  | 1000 | 0.19 | 0.15 | **0.04** | **0.001** | **0.001** |
|  | All | 0.1 | **0.03** | **0.05** | **<0.0001** | **0.006** |
| **Pyrazinamide** | 20 | **0.01** | 0.17 | **0.0005** | **0.01** | **0.01** |
|  | 50 | **<0.0001** | 0.08 | **0.0003** | **<0.0001** | **<0.0001** |
|  | 100 | **0.005** | 0.19 | **0.0008** | **<0.0001** | **<0.0001** |
|  | 500 | **0.05** | 0.06 | **0.0002** | **<0.0001** | **<0.0001** |
|  | 1000 | **0.04** | 0.06 | **0.0002** | **<0.0001** | **<0.0001** |
|  | All | 0.3 | 0.3 | **0.01** | **0.007** | 0.3 |

**Supp. Figure 1. Prediction results for resistance to Rifampicin**

Distribution of prediction error for resistance to rifampicin due to mutations in *rpoB* in the global dataset. A, B, C, D, E show subsampling results, whereas F shows the box plot for predicting resistance using the whole training set on the 20 random test sets.

**Supp. Figure 2. Prediction results for resistance to Isoniazid**

Distribution of prediction error for resistance to Isoniazid due to mutations in *katG* in the global dataset. A, B, C, D, E show subsampling results, whereas F shows the box plot for predicting resistance using the whole training set on the 20 random test sets.

**Supp. Figure 3. Prediction results for resistance to Pyrazinamide**

Distribution of prediction error for resistance to Pyrazinamide due to mutations in *pncA* in the global dataset. A, B, C, D, E show subsampling results, whereas F shows the box plot for predicting resistance using the whole training set on the 20 random test sets.

**Supp. Figure 4. Comparing range-normalized MetaDistance to MetaDistance**

Performance of range-normalized MetaDistance to predict drug resistance to Pyrazinamide due to mtuations in *pncA*. A shows the performance on the whole training set, and B shows subsampling results.
