## Supplementary figures and images for "A composite method to infer drug resistance with mixed genomic data"

### Supplemental Figure 1

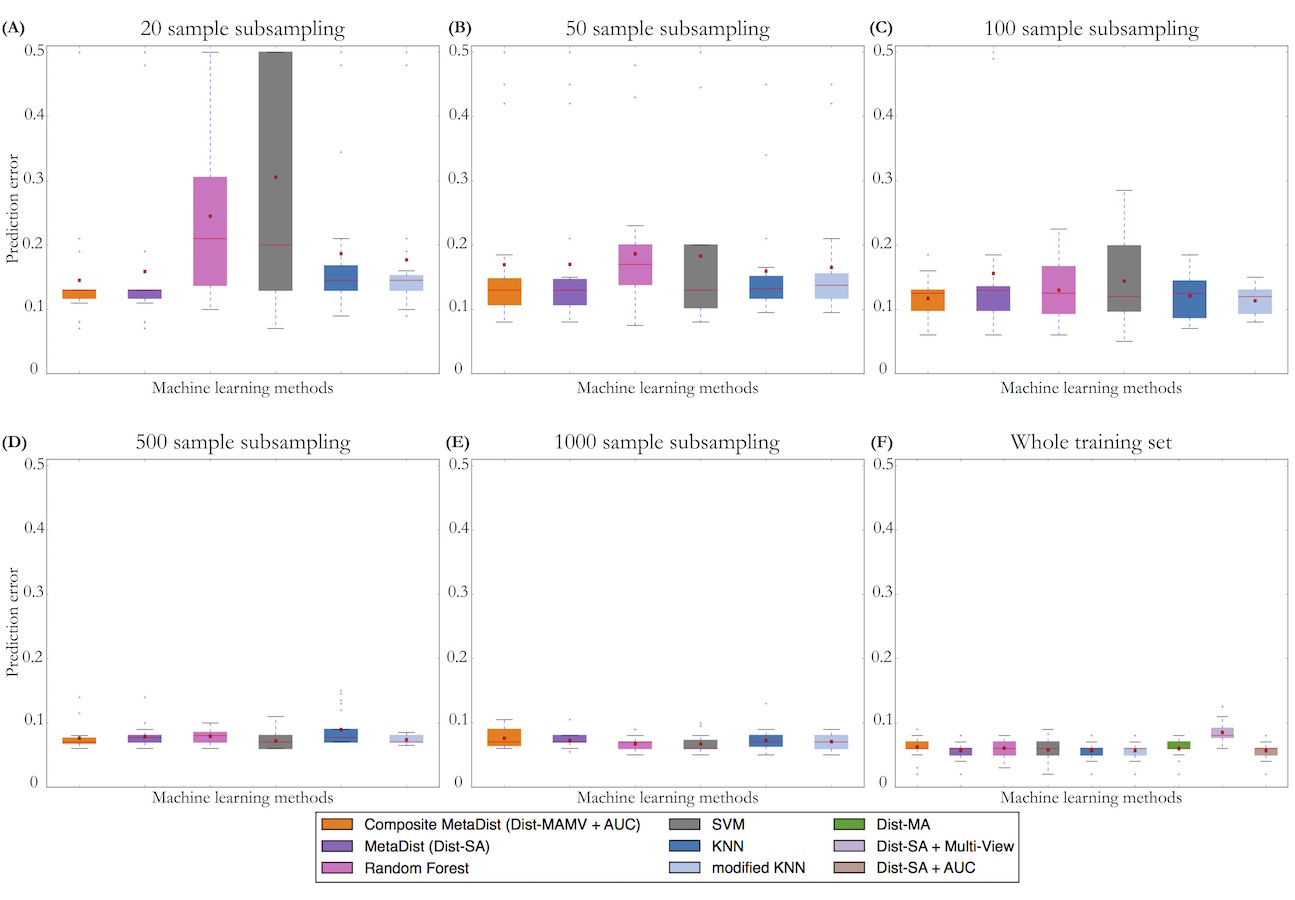

### Supplemental Figure 2

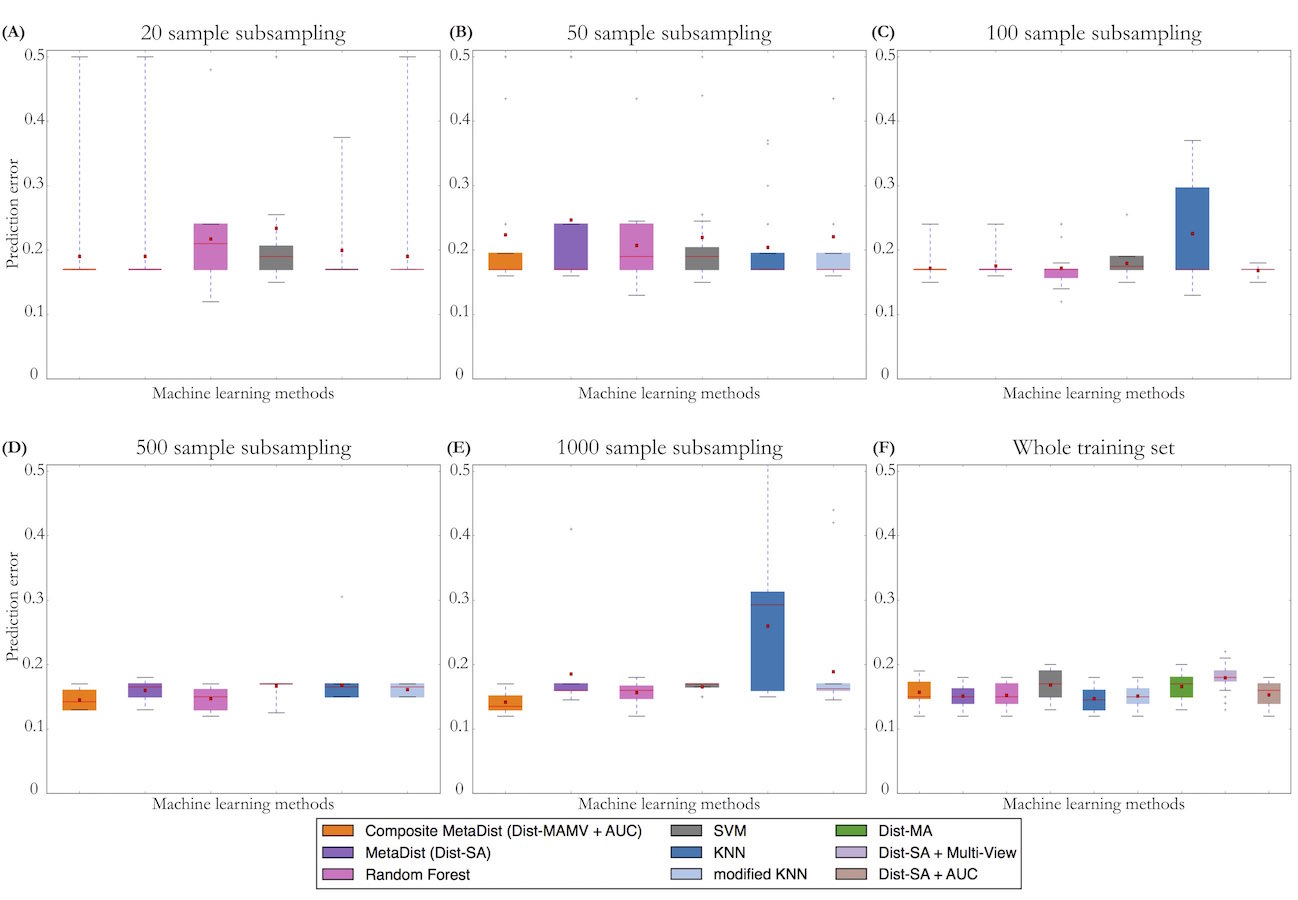

### Supplemental Figure 3

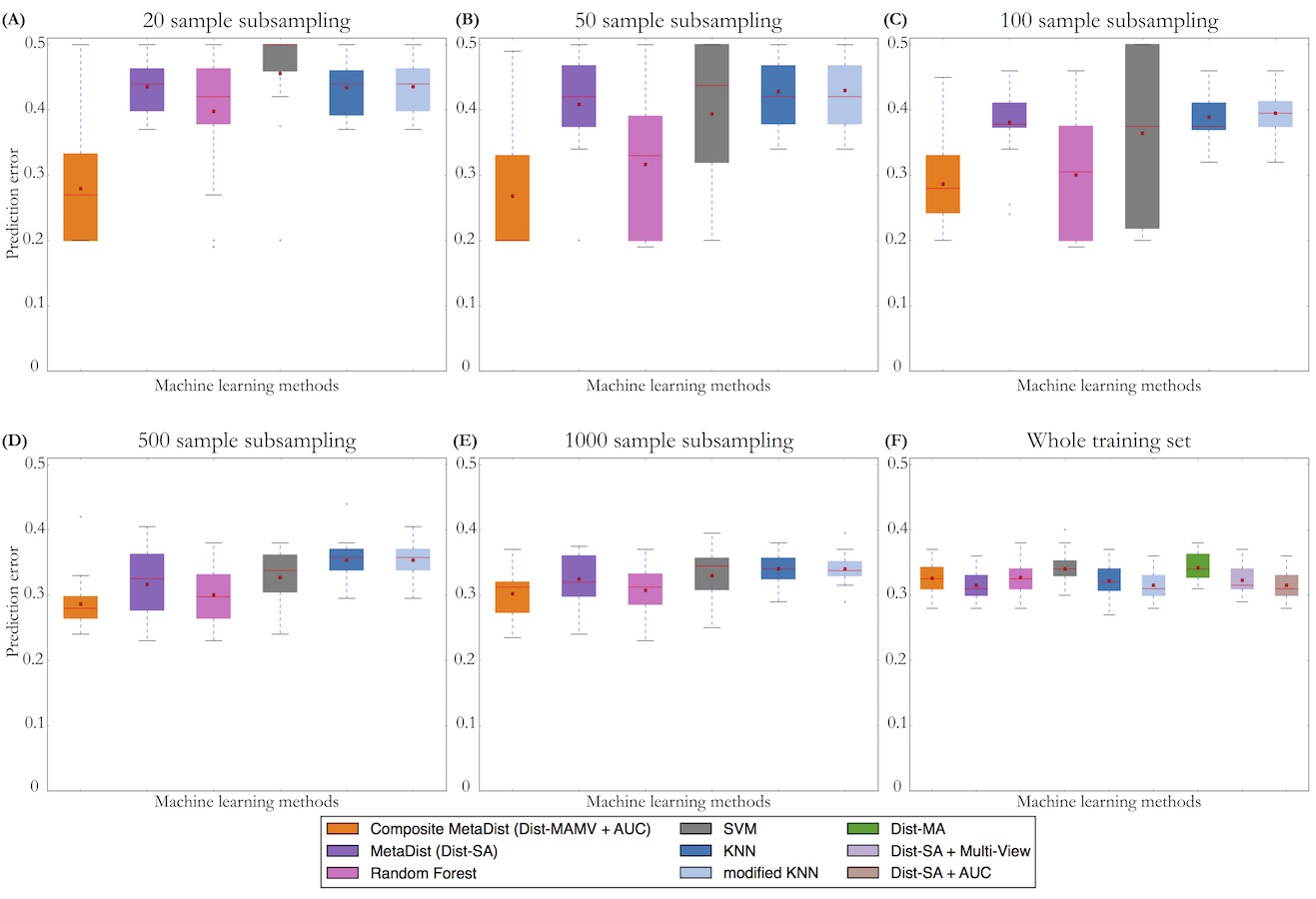

### Supplemental Figure 4

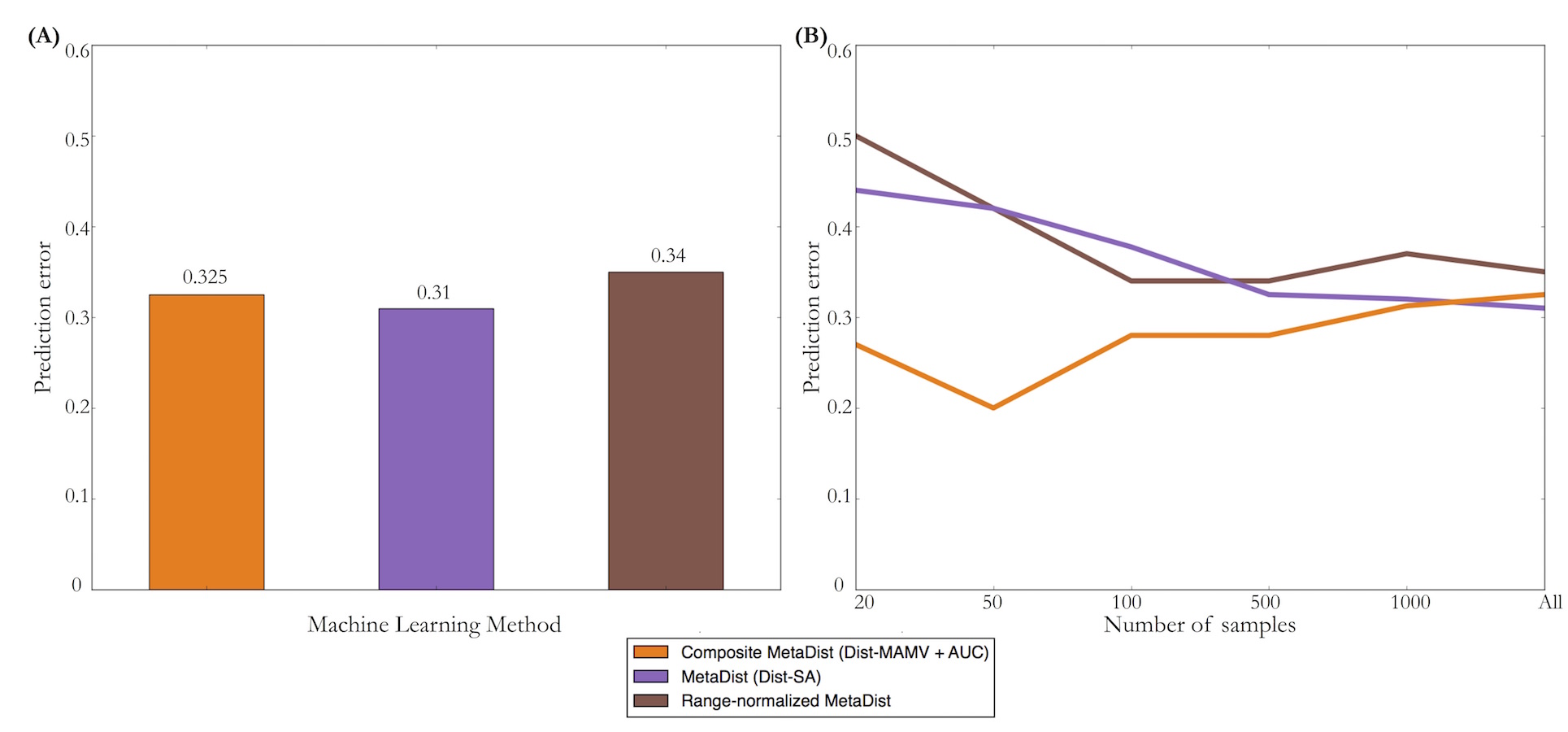
